# Metabolic heat flow from the minimal cell JCVI-Syn3B reveals the lipidome-dependence of growth and metabolism

**DOI:** 10.1101/2025.09.12.675810

**Authors:** Jana Oertel, Nataliya Safronova, James Sáenz, Karim Fahmy

**Author notes:** Universitätsklinikum Jena, Am Klinikum 1, 07747 Jena, Germany (Jana Oertel). Max Planck Institute for Terrestrial Microbiology, Karl-von-Frisch-Strasse 10, 35043 Marburg, Germany (Nataliya Safronova). Corresponding authors: Karim Fahmy, James Saenz.

## Abstract

The cell membrane facilitates interactions with the environment and serves as an organizational platform for coordinating cellular processes, with lipids playing a central role in determining membrane property and function. Yet, how lipidome composition influences cellular fitness remains poorly defined. Recent approaches to chemically tune and minimize the lipidomes of genomically minimized bacterial organisms such as JCVI-Syn3A/B offer a streamlined system to explore why cells need such diverse lipid chemistries. In this study, we use isothermal microcalorimetry to assess how changes in lipid composition affect heat dissipated by JCVI-Syn3B cells, a parameter reflecting both growth and metabolic efficiency. By transposing the Monod equation into a calorimetric equation and extending it to the full life times of batch cultures, we introduce a new approach to quantify the metabolic efficiency of JCVI-Syn3B. Remarkably, our results demonstrate that tuning lipidome composition results in considerable variations of energy dissipation at the expense of biomass production. As a consequence, the volume of these minimal cells becomes inversely coupled to the lipidome-dependent entropic cost of cell division. The corresponding change in heat flow per cell mass gives rise to a complex but systematic dependence of growth rates on lipid composition. Interestingly, the maximal rate correlates with maximal lipid diversity, suggesting that the ability to tune local cell membrane charge and curvature through lipid structural diversity is crucial for divisome function. Our observations highlight the critical role of lipidome composition in cell metabolism and growth, and provide a new tool for interrogating the relationship between membrane composition and cell fitness.

## Introduction

The complexity of the lipid composition in biological membranes contrasts the simplicity of its prime physical function, namely, the maintenance of a diffusion barrier between the cell and its environment. Whereas a barrier function can be fulfilled by a single lipid species, lipid structural diversity must have evolved for other vital membrane-associated processes. For example, membrane protein function has been investigated intensely *in vitro* using various dedicated membrane-mimetic systems with controllable lipid composition as reviewed.(1–3) Besides specific lipid protein interactions, global membrane properties such as bilayer lateral pressure(4, 5) lipid asymmetry(6) as well as membrane curvature(7, 8) tune the conformation and functional transitions of membrane proteins and of membrane-associated processes, including cytoskeletal interactions.(9) Thus, a variety of lipidome-related processes may affect cell biochemistry through altered substrate or ion transport efficiencies or perturbed plasma membrane mechanics and may ultimately affect metabolic activity through impaired energy utilization.

Lipid-specific as well as ensemble properties of lipids in the plasma membrane will eventually couple to metabolic processes by affecting transmembrane transport, osmoregulation, signaling, cytoskeletal activity a.o.. Such impact on cellular functions has been studied mostly in eukaryotic systems (10) but has been recognized in bacterial cells as well(11–13) and can be considered a fundamental aspect of life. However, gaining control over the lipid composition in a native plasma membrane, while preserving intracellular biochemistry, represents a crucial conceptual challenge in studying the interplay between lipid complexity and cell biology in a living cell. The mere provision of a defined lipid mixture to cultured cells results in much higher lipid diversity than of that provided in the diet, since lipids become remodeled by the enzymatic machinery of the cell.(14) This metabolic interference has been largely circumvented by the use of genome-minimized cells derived from *Mycoplasma mycoides*.(15, 16) These “minimal cells” retain only a very limited capability to synthesize lipids and thus rely critically on lipids taken up from the growth medium, but still show lipidome adaptation to their environment.(17) Recently approaches for chemically tuning and minimizing the lipidome of the “minimal cells” JCVI-Syn3A and JCVI-Syn3A/B (nearly identical strains, referred to collectively here as JCVI-Syn3A/B) have been introduced, providing an opportunity to study the effect of lipidome complexity on growth and metabolism of a biochemically streamlined system.(14, 18) What remains unknown is how the membrane composition of a cell with an enzymatic network of reduced complexity would affect the distribution of energy flux between direct biomass-forming metabolic activities and other energy-consuming processes such as lipid remodeling or cell division that may depend critically on lipid complexity.

We surmised that the minimal adaptability of JCVI-Syn3A/B may facilitate a systemic approach to lipid-related metabolic cost estimates by monitoring the amount of energy dissipated as heat during the growth of batch cultures of “minimal cells”. The rationale behind this approach is the suspicion that the restricted adaptability of plasma membrane properties to vital cellular functions will render metabolism less efficient for growth and cell replication due to “extra” heat dissipation from metabolic activities that would normally require optimally tuned plasma membrane properties. The obvious challenge of our endeavor is the lack of chemical specificity in metabolic heat flow measurements. In fact, metabolic heat represents probably the least specific signal an organism can offer, because it originates in the sum of all of its net exothermic biochemistry. On the other hand, calorimetric measurements have become extremely sensitive and provide the enormous advantage that any change in cell biochemistry can in principle be captured by such experiments when compared to a suitable reference.(19–24)

Here, we have investigated the influence of seven defined lipid diets on the growth of JCVI-Syn3B cells. A comprehensive analysis of differences in the corresponding metabolic heat flow signatures measured by isothermal microcalorimetry (IMC) was performed to derive previously unknown growth parameters of JCVI-Syn3B cells such as accurate growth rates and relative measures of per cell metabolic heat production. These data carry information on the energetic expenditures that support cell replication as a function of plasma membrane lipid composition. We suspected that it should be possible to detect even small lipid-specific enthalpic changes in cell metabolism on the background of a shared cell biochemistry in all diets driven by glycolysis (JCVI-Syn3B lacks a TCA cycle) which provides the prime carbon-based metabolic building blocks and the sole energy source. In this regard, we found surprisingly large variations in the thermal signatures of minimal cells grown in various lipid diets.

In order to assess the potential wealth of information from these metabolic heat flow data, we developed new quantitative analysis tools based on an extended calorimetric form of the Monod equation. By combining calorimetric data with lipidomics and cell size determinations, we obtained a comprehensive per cell view of how lipid composition affects nutrient turnover, cell division rate and biomass yield. Surprisingly, the energetic cost of biomass formation and the rate of cell division were differently affected by the diets. This trade off conforms with unexpectedly systematic relations between cell volume, metabolic heat flow per cell mass and division rate in response to lipid complexity.

## Results

### Tuning the lipidome composition in JCVI-Syn3B

We previously demonstrated that *M. mycoides*, JCVI-Syn3A/B can take up exogenous free fatty acids to synthesize phosphatidylglycerol (PG), Cardiolipin (CL) but also low abundances of phosphatidic acid (PA) and sometimes diacylglycerol (DAG).(14, 18) To support growth, cells require at least one saturated and one unsaturated fatty acid, typically C18:1 (oleate) and C16:0 (palmitate). This diet, designated 2FAs, imposes a constraint on phospholipid remodeling: acyl chain unsaturation and length are coupled and cannot be independently remodeled. To lift this constraint, we designed a diet with two additional fatty acids, C18:0 and C16:1. This more diverse lipid diet, designated 4FAs allows independent variation of phospholipid chain length and unsaturation, increasing degrees of freedom available to the lipidome, allowing us to observe the effect of increased acyl chain diversity.

JCVI-Syn3A/B can also take up exogenous phospholipids, providing a means to tune lipid class diversity. We analyzed the lipidome of cells grown on a rich lipid diet provided from fetal bovine serum (FBS).(17) This work showed that in addition to cholesterol, phosphatidylcholine (POPC) and sphingomyelin (SM) were major components of the lipidome. Therefore, to assess the role of lipid class diversity, we extended the 2FAs diet by sphingomyelin (2FAs_SM), or by POPC designated 2FAs_PC, or by bothlipids, designated 2FAs_PC_SM. JCVI-Syn3A/B requires a sterol for growth, so all diets contained cholesterol. Lipidomic composition for cells grown on these defined lipid diets are reported separately in Safronova et al..(18)

### The metabolic activity of JCVI-Syn3B cells follows Monod kinetics

We have studied by isothermal microcalorimetry (IMC) the metabolic heat flow from minimal cells of the strain JCVI-Syn3B cultured in seven lipid diets of increasing complexity. For each diet, IMC data were recorded in triplicate (Fig. S1) and evaluated individually as exemplified in Fig. 1 for the two diets 2FAs and 2FAs_SM that produced the largest and the smallest heat release among all tested growth conditions, respectively. IMC data were acquired as thermal power (heat flow) *vs*. time *t* (Fig. 1A) expressed in most general terms as

**Figure 1:**
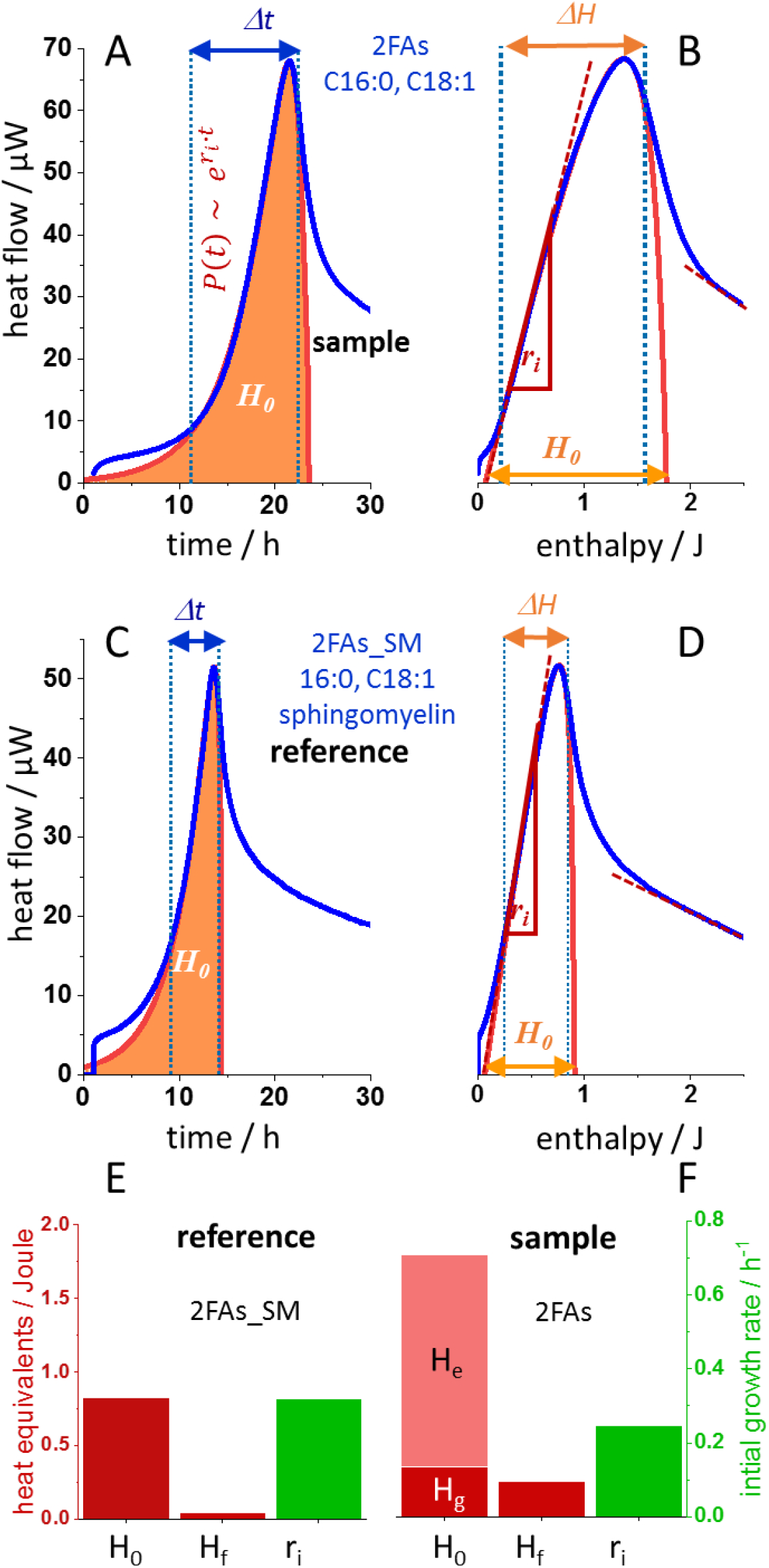
Heat dissipation from JCVI-syn3B cells varies in a lipid-dependent manner. (A) Heat flow from cells grown on a lipid diet with cholesterol and two fatty acids(2FAs: palmitic & oleic acid). After baseline stabilization, the heat flow increased exponentially with rate *r*_*i*_. Dotted lines (blue) delimit the growth phase following the ECME, (Eq 2). Fits (red) can be extrapolated “back” to early exponential growth and “forward” to full nutrient consumption, where total heat *H*_*0*_ equals the integral over the fitted curve (orange). (B) The same data plotted *vs* the released heat *H* (exponential phase corresponds to straight line of slope *r*_*i*_). (C, D) the same evaluation of IMC data from cells grown in the 2FAs diet supplemented with sphingomyelin (2FAs_SM). (E,F) Parameters *H*_*0*_, *H*_*f*_ and *r*_*i*_ obtained for growth in the two diets. *H*_*0*_ from the reference provides a heat equivalent *(H*_*g*_) for biomass (dark red) as opposed to additional “extra” metabolic heat *H*_*e*_.

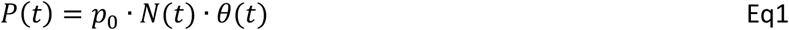

with *p*_*0*_, the maximal metabolic heat flow per cell; *N*, the number of cells and *θ*, the “metabolic load”. The latter is a unitless number between zero and one expressing the actual heat flow per cell as a fraction of *p*_*0*_. We described the “metabolic load” *θ* by the Monod equation (ME) by transforming the data into a function of enthalpy *H* (released heat). *H* scales with the amount of <u>consumed</u> nutrient, such that *H*_*0*_*-H* can replace the substrate concentration in the original ME leading to:

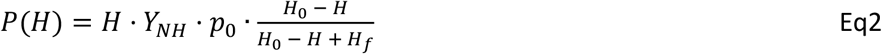

with *Y*_*NH*_, the number of cells formed per Joule of released metabolic heat. The maximal growth rate of the cells at nutrient saturation is *r*_0_ = *Y*_*NH*_ · *p*_0_. The fraction on the right side expresses *θ* in the form of a “calorimetric” ME, where *H*_*f*_ is the mathematical homolog of the Monod constant (*C*_*M*_). This “extended calorimetric Monod equation” (ECME) differs from the original ME by the additional leading term *H*. The latter accounts for the fact that the number of metabolizing cells in a growing culture increases in proportion to the amount of released heat. *P(H)* follows an inverted parabola which can be fitted to the IMC data according to Eq2. This was done in the maximal heat interval *ΔH* that complied with the Monod-type of growth exempliied in Fig. 1 A and B for the 2FAs diet in the time and the enthalpy domain, respectively. More than 80% of a Monod-type of growth was covered by the IMC data of JCVI-Syn3B cells in all seven tested lipid diets. Beyond that range, data where not analyzed. The fit for the minimal 2FAs diet predicts a maximal cumulative heat release of *H*_*0*_ = 1.77 J if the culture had grown to its final biomass in compliance with the ECME. Figures 1 (C,D) show the corresponding results for JCVI-Syn3B cells grown in the additionally presence of sphingomyelin (2FAs_SM). Here, growth was accelerated at smaller heat flow and reduced heat release of *H*_*0*_ = *0*.*83 J*. Thus, with the identical amount of nutrients, “minimal cells” in the 2FAs diet grew about half as efficiently as in the reference diet.

### Metabolic heat flow is strongly coupled to plasma membrane composition

How can the lipid-specific energy expenditure be differentiated from “normal growth-related” energy flux? In the 2Fas_SM diet, “minimal cells” produced the smallest amount of heat, rendering it a reference to rank energy efficiency for all other cultures. Due to the “low complexity” of JCVI-Syn3B cells, metabolic adaptation in response to the lipid diet is limited, if present a all, leaving the majority of cell biochemistry and <u>molar</u> reaction enthalpies unaffected. Thus, a fraction *H*_*g*_ *= Y*_*g*_ *· H*_*0*_ from the total heat release from all cultures can be assigned to these reactions, for which *Y*_*g*_ expresses a yield in a non-mechanistic manner with *Y*_*g*_ *= 1* for the reference culture.

Following this heuristic approach, we modeled the culture growth in the 2FAs diet with the parameter *H*_*f*_ from the 2FAs_SM reference, assuming that only a fraction *α* of the metabolic flux of nutrients was used for “reference-like” growth in the 2FAs diet. The remaining fraction *(1-α)* was assigned to lipidome-dependent processes (Eq S1). The mechanistic parameter *α* replaces the descriptive yield *Y*_*g*_ such that *H*_*g*_ = *α* · *H*_0_ and a lipidome-dependent “extra metabolic heat” can be defined as as *H*_*e*_ = (1 − *α*) · *H*_0_. Only about 20% of the total heat release of the culture in the 2FAs are predicted to be related to the shared cell biochemistry which produces less than half the biomass of the reference culture (Figure 1 (E, F)). Surprisingly, the initial growth rates *r*_*i*_ (the initial slope of the enthalpy plots) were much less affected. If indeed *H*_g_ provides a measure of biomass, the data imply that the change in the plasma membrane lipid composition affected also cell size, because at comparable division rates, less biomass formed in the 2FAs diet would be partitioned among a similar number of cells as in the reference diet.

### Lipidome complexity controls the balance between cell division rate and biomass formation

The bar graph in Figure 2A shows the parsing of total metabolic heat *H*_*0*_ for cultures grown in the tested lipid diets in the order of increasing growth efficiency (decreasing *H*_*0*_). In the same order also the biomass proxy *H*_*g*_ and the “extra metabolic” heat *H*_*e*_ exhibit a monotonic increase and decrease, respectively, indicative of biomass being sacrificed for lipidome-dependent biochemistry. The maximal growth rates *r*_*0*_ are shown as well. In contrast to the initial growth rates, they are independent of the initial nutrient concentration. The exemplified larger effect of the lipid diets on biomass than on *r*_*i*_ also holds for the maximal growth rates *r*_*0*_ in all diets. The implied alteration of cell size was tested by dynamic light scattering (DLS) of the cells as shown in Fig. 2 (insert box) and is remarkably consistent with the ranking by *H*_*g*_. However, the maximal heat flow per cell normalized by the total heat release and by cell surface a (*p*_*0na*_ *= p*_*0*_*/(H*_*0*_*·a*)) did not vary much. In other words, the cell mass was strongly affected by the lipid composition but not the flux of nutrients per cell surface (Table S1).

**Figure 2:**
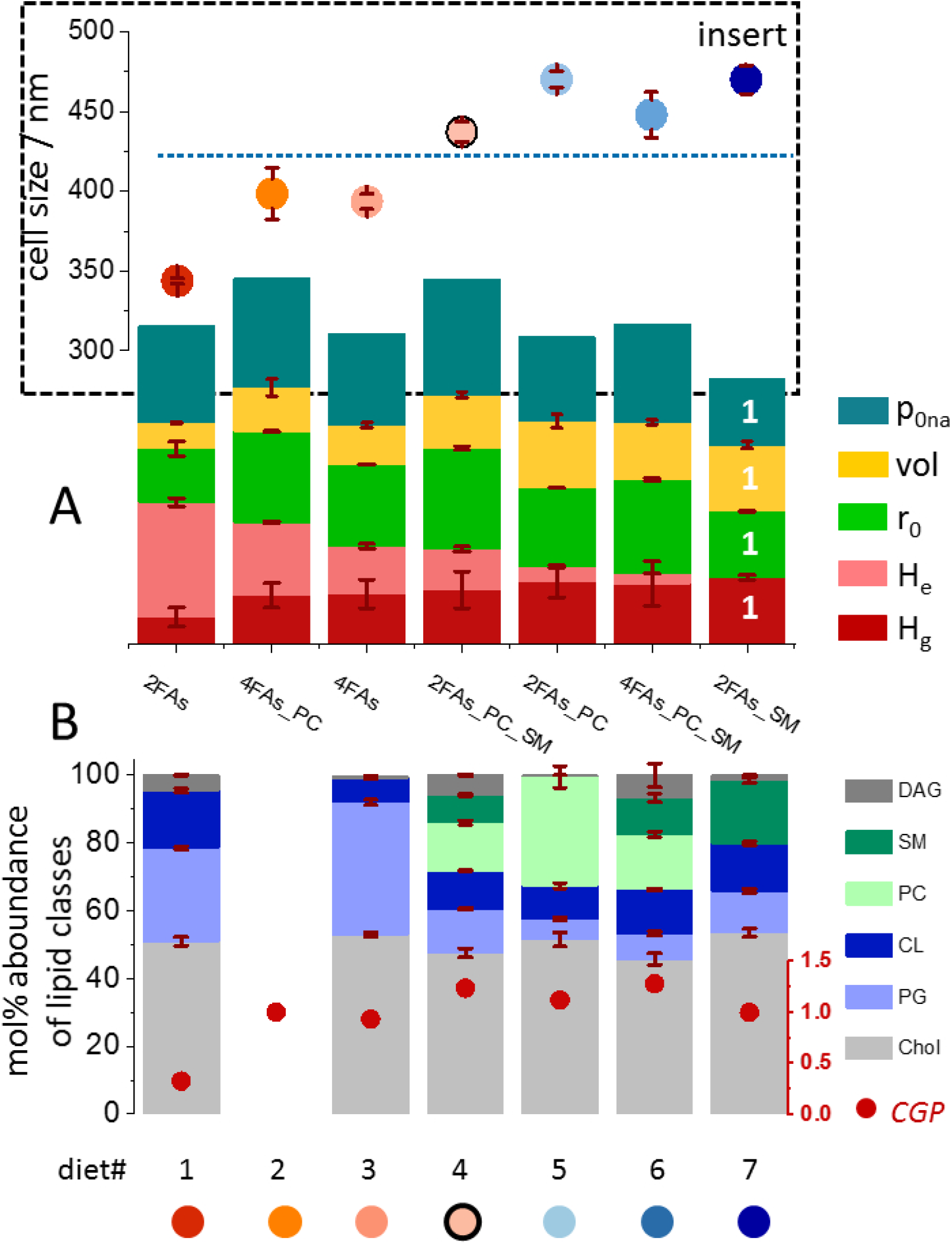
Membrane lipid composition influences the balance between biomass production, growth rate, and metabolic efficiency. Upper bar graph. *H*_*g*_ (dark red, biomass proxy) fraction of total heat released in accordance with the Monod parameters of the diet 2FAs_SM); *H*_*e*_ (light red), lipidome-dependent extra-metabolic heat; *r*_*0*_ (green), maximal cell division rate; *vol* (yellow), relative cell volume; *P*_*0na*_ (turquoise), relative nutrient flux (*p*_*0*_*/H*_*0*_) normalized to cell surface area. Parameters are normalized with respect to their values in the reference diet (2FAs_SM, set to unity). Lower bar graph: abundances of lipid classes (in mol% of total lipid) in plasma membranes from JCVI-syn3B cells grown on defined lipid diets. DAG: diacyl-glycerol; SM: sphingomyelin; PC: phosphatidyl-choline; phosphatidyl-glycerol; Col: cholesterol. Culture growth performance *CGP* = *r*_*0*_ *· H*_*g*_ (red filled circles), Insert Box: cell diameter determined by dynamic light scattering for each lipid diet. Number values of the data are given in Table S1. Data from different diets are color-coded red to blue, from large to small heat dissipation.

Aiming at correlating kinetic and calorimetric results with theplasma membrane lipidome, JCVI-Syn3B cells were subjected to lipid class analysis (Fig. 2B), covering the least and the most efficient biomass-forming cultures as well as four cultures with intermediate heat releases. Furthermore, we have included a descriptive <u>C</u>ulture <u>G</u>rowth <u>P</u>erformance parameter *CGP = r*_*0*_ *· H*_*g*_ (normalized with respect to the reference culture). It expresses the maximal rate of biomass formation and shows that the cells performed the worst in the 2FAs diet. The addition of sphingomyelin (FAs_SM diet) increased the biomass but with little effect on division rate, whereas the *CGP* was highest in the most complex diets, where POPC predominantly enhanced division rate.

### The plasma membrane lipidome determines the entropic cost of JCVI-Syn3B cell division

The heuristic dissection of metabolic heat into *H*_*g*_ and *H*_*e*_ based on different substrate utilization ratios in the ME has revealed key features of the lipidome-dependent energy expenditure: cell biomass is sacrificed for lipidome-dependent processes with unexpectedly large energy dissipation. Since *H*_*g*_ scaled with cell volume, it is a suitable proxy of biomass formation (Fig. 3A). However, cells in the reference culture also perform all necessary non-biomass-related metabolism. In order to differentiate enthalpy contributions independently of a reference culture, we denote in the following the “true biomass-related heat” as *H*_+_ and the “true non-biomass-related heat” *H*_−_. By its definition, the “true biomass yield” *Y*_*BH*_ follows a logistic curve in dependence on *ln(H*_+_*/H*_−_) (see SI).

**Figure 3:**
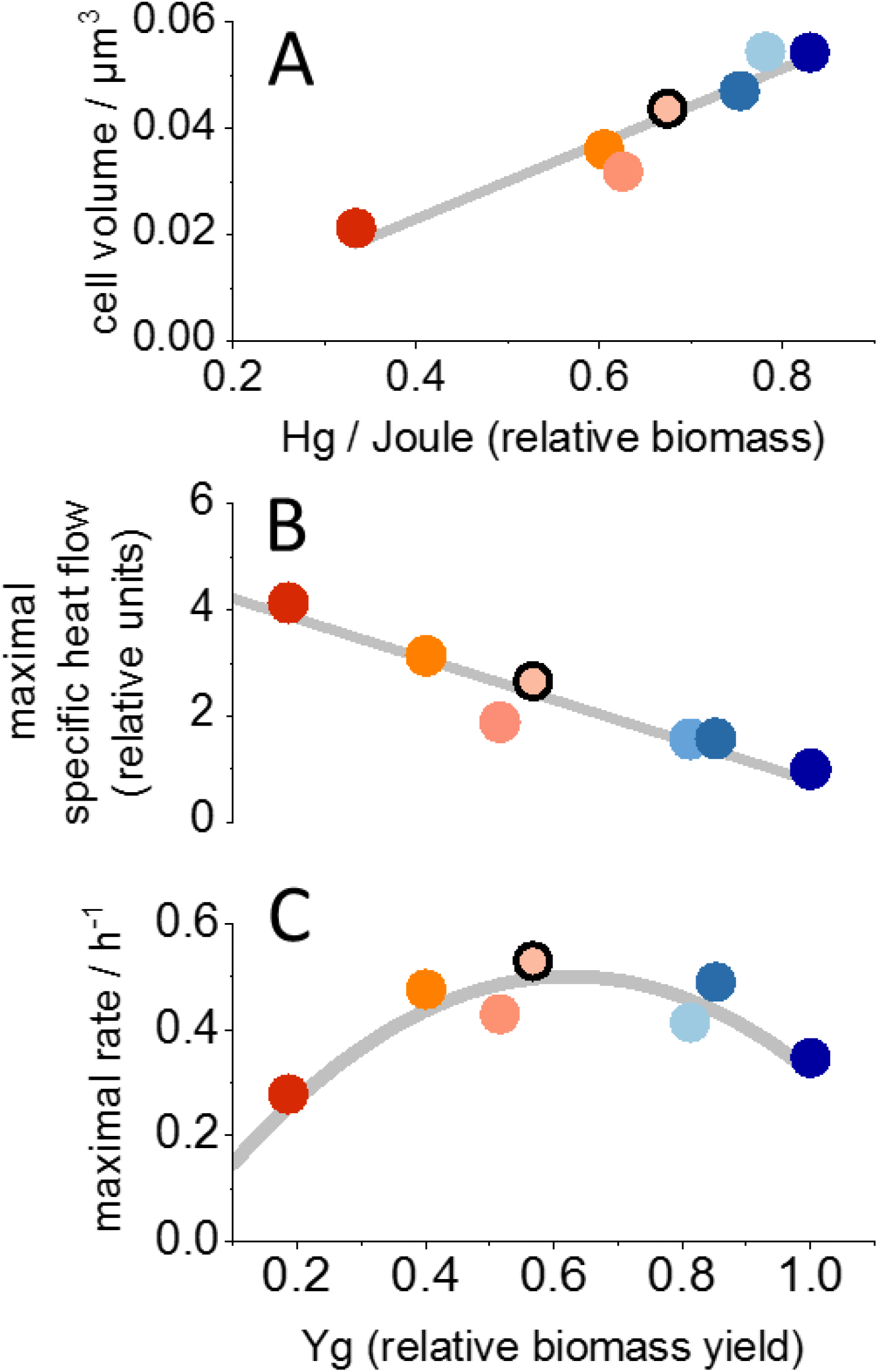
Calorimetry-derived energy partitioning indicates a tradeoff between cell division rate and biomass yield. (A) Proportionality between the calorimetric biomass measure *H*_*g*_ and the cell volume. (B) The “maximal specific heat flow” *p*_*0m*_=*p*_*0*_*/m* at substrate saturation declines linearly with biomass yield: *p*_*0m*_=*P*_*0m*_ *· (1-Y*_*g*_). (C) Parabolic dependence of maximal cell division rates on the biomass yield *Y*_*g*_. Color code as in Fig. 2.

How is the “true biomass yield” *Y*_*BH*_ related to the fitness of JCVI-Syn3B cells? The reference-dependent *CGP* values in Fig. 2B provide a measure of the rate of biomass formation and show that high lipid complexity supported “fitness”. The latter is given by the product of cell mass *m*_*c*_ and maximal division rate *r*_*0*_ which can also be expressed as the product of the calorimetric parameters *p*_*0*_ and *Y*_*BH*_ leading to the following equality:

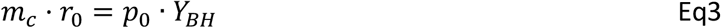

This relation is key to understanding the intriguing response of cell division rates to plasma membrane lipid composition, because it shows that the maximal division rates depend on the maximal “specific heat flow” *p*_*0m*_ *= p*_*0*_ */ m*_*c*_.

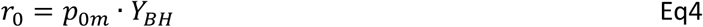

Our data show that the cell mass-specific heat flow *p*_*0m*_ is not constant but itself a function of biomass yield (Fig. 3B) leading to a parabola-shaped dependence of *r*_*0*_ on *Y*_*BH*_ and thus also on *Y*_*g*_ (Fig.3C). The transformation of *Y*_*g*_ into the reference-independent yield *Y*_*BH*_ is described in the SI (Eq S8/8a). With the appropriate scaling of the *H*_*g*_ values, the “true biomass” yield follows indeed the expected logistic curve. In the same representation, the *r*_*0*_ values are well reproduced by Eq4 with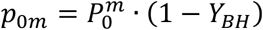, where 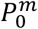 formally defines an upper limit of the “maximal specific heat flow” of a cell in an “optimal diet”.

Evidently, the calorimetric data obey well-defined quantitative relations. Whereas the true biomass yields (normalized from zero to one) could be obtained by mere scaling of the biomass proxy *H*_*g*_, the cell division rates exhibit a non-intuitive lipidome-dependence. In order to describe metabolic fitness based on the enthalpy contributions *H*_+_ and *H*_−_, we introduce the corresponding pathway-specific yield parameters *β* and *δ* which relate biomass formation and cell division rate to the reaction enthalpy of the underlying biomass- and non-biomass-forming biochemistry, respectively. The corresponding equations *r*_*0*_ = *δ* · *H*_−_ and *m* = *β · H*_+_ allow reconstructing the complete “fitness landscape” of biomass production rates as a function of the partial enthalpies *H*_+_ and *H*_−_ (Fig. 4B). The experimentally obtained rates of biomass formation cut through this “fitness landscape”, tracing out one out of many possible trajectories in the 2D-plot. Remarkably, the linear relation between *H*_+_ and *H*_−_ shows that the ratio of the pathway-specific differential reaction enthalpies stayed roughly constant (*ΔH*_+_*/ΔH*_−_ ~-0.3), i.e., the heat released from non-biomass reactions increased in proportion to the degree by which biomass was sacrificed. The cross-cut section through the “fitness landscape” reproduces the one-dimensional presentation of fitness based on the *CGP* values in Fig.2B. The fitness landscape was generated with a diet-independent pathway-specific yield *β* for biomass synthesis in agreement with the experimentally obtained good correlation between *H*_*g*_ and cell volume in all diets. In contrast, the yield *δ* obtained from the experimental *r*_*0*_ values (Eq S12a)) depended on the lipid diet and decreased with cell mass (Fig. 4C): the more heat was released to support a division rate of 1 h^−1^, the less biomass was formed.

**Figure 4:**
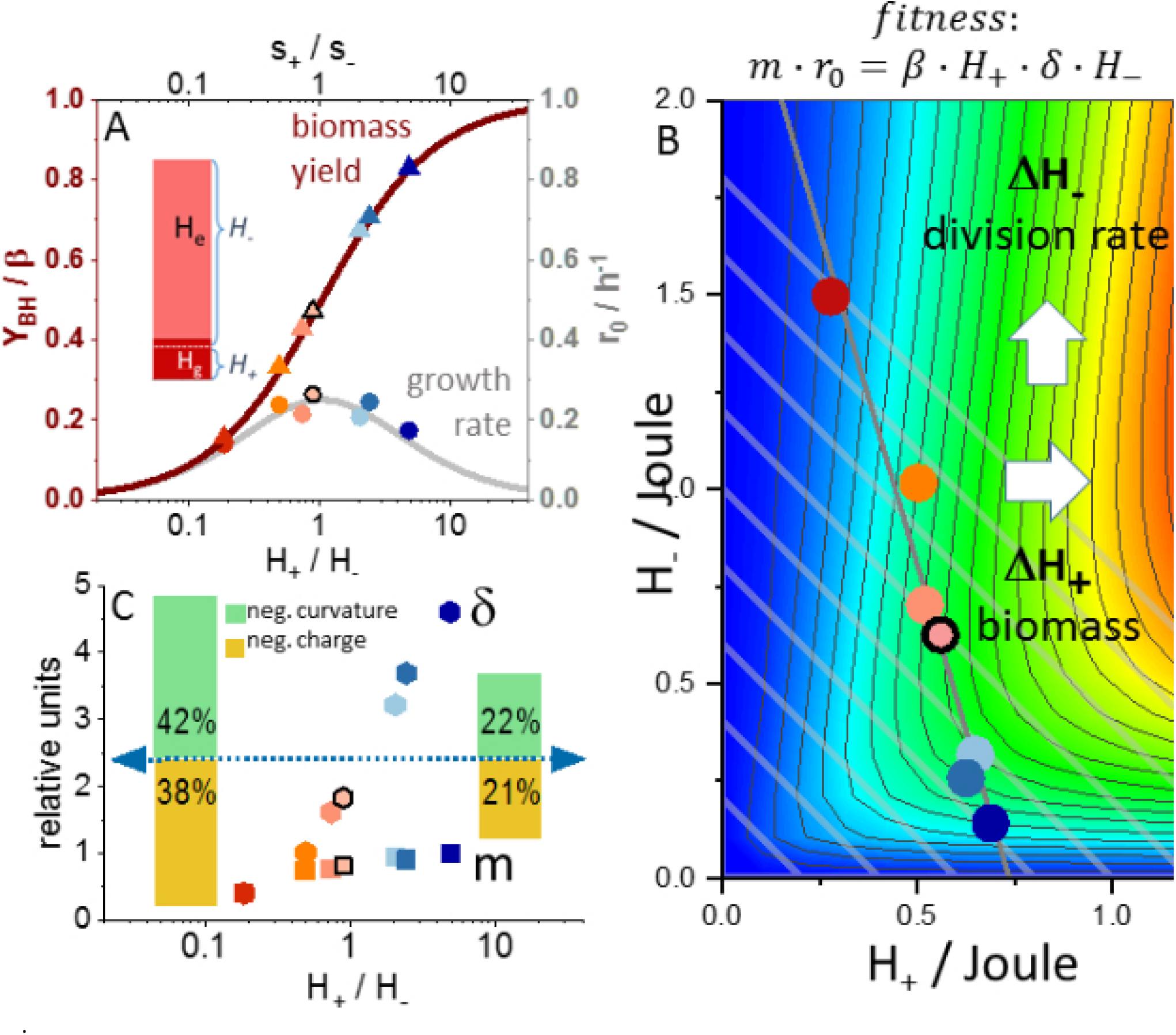
Energy partitioning is coupled to biomass yield, growth rate, and cellular fitness. (A) Normalized biomass yields *Y*_*BH*_ (triangles) plotted against *ln(H*_+_*/H*_−_) follow the predicted sigmoid (Eq S8a), when the biomass-related heat was expressed as *H*_+_ = 0.83·*H*_*g*_ (exemplified by the bar graph insert) to account for non-biomass contributions to *H*_*g*_. With this correction, the maximal division rates *r*_*0*_ follow a logistic distribution (grey). (B) “Fitness landscape” representing the rate of biomass formation *m*_*c*_*·r*_*0*_ as a function of biomass-(*H*_+_) and non-biomass-(*H*_−_) related heat components. We used a constant pathway-specific yield *β* for biomass production and a lipid-dependent yield *δ* for the division rate (Eq S12). “Isocaloric” lines (grey) correspond to constant tootal heat release *H*_*0*_ which can be read from the x- or y-axis interceptions. C) Relative yield *δ* for division rate (filled circles) and relative cell mass (filled squares) are positively correlated. Relative changes in lipid composition of the diets with respect to charge and intrinsic curvature are indicated by bars of the insert.

### Flux balance analysis reveals the lipidome-dependent rate and energy efficiency of cell division

The described quantitative relations allow reconstructing the calorimetric, kinetic and volumetric data in the form of an equivalent electronic circuit, equivalent to a flux balance analysis. Four resists are sufficient to construct such a circuit as shown in Fig. 5A. Flux through the constant resists *R*_*b*_ and *R*_*d*_, represent energy flowing into biomass production and cell division, respectively, whereas current through *R*_*f*_ corresponds to futile energy dissipation in the cell division machinery. The metabolic heat flow is represented by the cumulative currents through these resists with each current contributing heat with a constant weighing factor (not the electro-thermal energy generated by the resists themselves!). Over one doubling time *τ*, these currents *I*_+_ and *I*_−_ transport a charge in proportion to *H*_+_ and *H*_−_, respectively. The resist *R*_*a*_ determines the total amount of total flux *I*_−_ through the divisome, thereby accounting for lipidome-dependent alterations of energy barriers in functional transitions of the divisome. Lipid-dependent energy barriers do not leave a calorimetric signature, i.e., *I*_*a*_ has a weight of zero, but affect metabolic kinetics. The equations for *R*_*f*_ and *R*_*a*_ can be solved such that the current through *R*_*d*_ reproduces the logistic distribution of growth rates (Fig. 4A) as a function of *I*_+_*/I*_−_ (total current of 1 A). Thereby, the bar graph shown in Fig. 2A can not only be reproduced but also aligned along a meaningful continuous x-axis (Fig. 5B) which covers all possible ratios between biomass- and non-biomass-related fluxes (Fig. 5C). The simulation shows that the complex response of metabolism to lipidome alterations can be broken down in the impairment of cell division by both energetic inefficiency by adjusting *R*_*f*_ and slowed kinetics by adjusting *R*_*a*_. In the first case, an energy sink is produced which depletes the ATP pool for biomass production. In the second, biomass production proceeds longer between cell divisions without having experienced a change in its intrinsic rate or efficiency. The observed lipidome-dependent changes in cell volume are the necessary outcome of the competition of the two metabolic processes for energy resources, such that the relative cell volumes are obtained from the circuitry as the ratio *I*_+_*/I*_*d*_., which is simply the rate of biomass production divided by division rate.

**Figure 5:**
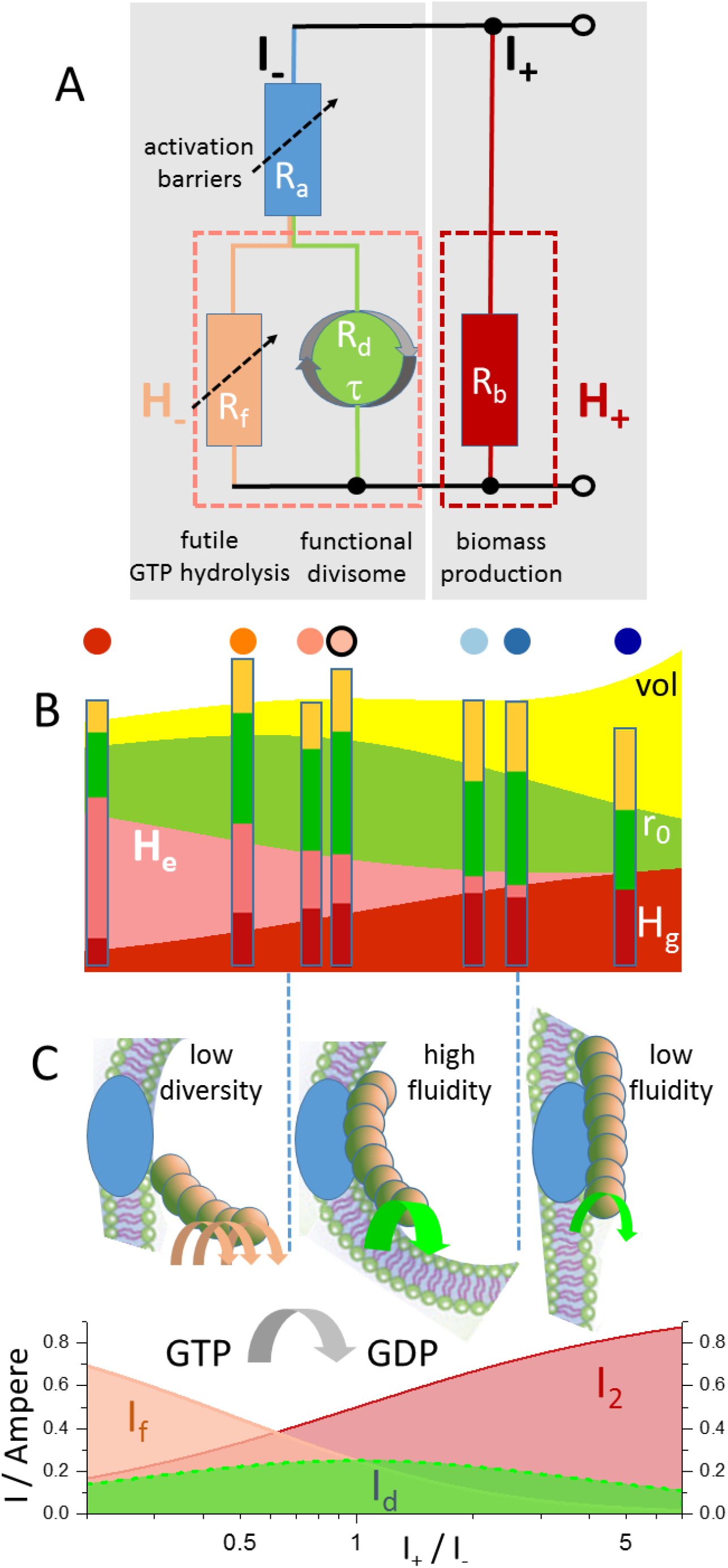
Flux balance simulation of lipidome-dependent metabolic energy fluxes in minimal cells. A) Equivalent electric circuit model. The currents *I*_−_ and *I*_+_ represent the fluxes of energy driving cell division (a.o. non-biomass-related processes) and biomass formation, respectively. The resist *R*_*a*_ attenuates the rate of cell division. The resists *R*_*d*_ and *R*_*f*_ determine the currents linked to energy flux in a functional divisome and futile GTP hydrolysis in an impaired divisome, respectively. Elements that contribute to heat flow in proportion to the current in each branch are boxed by dashed lines. B) Reproduction of the bar graph of calorimetric data shown in Fig. 2 as a function of log(*H*_+_*/H*_−_). C) Currents *I*_*d*_, *I*_*f*_ and *I*_+_ scale with divisome activity, futile GTP hydrolysis and biomass formation, respectively, as a function of log(*I*_+_/*I*_−_). The most representative lipid properties are indicated and non-productive (red arrows), productive (bold green arrow) and slowed productive divisome activities (green arrow) related to putative membrane protein interactions in the respective sketches (FtsZ: brown spheres, divisome-anchoring complexes: blue).

## Discussion

JCVI-Syn3A/B cells harbor the smallest life-supporting enzymatic network, making them a powerful model system to understand how cellular organization, biomolecular composition, and metabolism are mechanistically intertwined.(25) Fundamental benchmarks of the energetic and dynamic performance of their metabolism may thus be derived from such “minimal cells”. Given the complexity, variability and adaptability of the lipid composition of cell membranes in organisms spanning the domains of life, we wanted to know whether and how the lipid composition of the plasma membrane can couple to a rather primordial glycolysis-based metabolism. To answer this question, we have performed a comprehensive quantitative analysis of the metabolic activity of JCVI-Syn3B cells in which the lipidome can be tuned by chemically defined lipid diets provided in the growth medium. (14) In the formalism of an “extended calorimetric Monod equation” (ECME), metabolic heat flow curves from minimal cells showed an unexpectedly strong effect of the lipid diet on overall metabolic heat production. Changes in the lipidome were found to reduce cell biomass production by more than a factor of two and at the same time increased metabolic energy dissipation into heat. The limited capability of fatty acid modification (phosphorylation, glycerol addition and CL formation) and the small number of involved chemical bonds in relation to biomass-forming reactions exclude energy expenditure for lipid remodeling as the cause of the large biomass loss of JCVI-Syn3B cells grown in the presence of only two fatty acids (2FAs). Furthermore, PG formation was enhanced in the 4FAs diet at the expense of CL. Despite the higher demand for C3 metabolites per fatty acid chain in PG, biomass production was more efficient in the 4FAs diet. Likewise, the influx of nutrient cannot explain the observed changes in rates and yields. It was actually slightly larger in all lipid diets relative to the reference, varied less than the other growth parameters and did not correlate with any of these (Fig. 2). We conclude that plasma membrane composition influences metabolic efficiency not through lipid remodeling costs or impaired nutrient uptake, but by redirecting energy flows between biomass production and other cellular processes.

To quantify these lipidome-dependent energy fluxes, we separated the measured heat dissipation into a biomass-related *H*_+_ and a non-biomass-related component *H*_−_, both extrapolated to full substrate consumption (EqS8a). We further introduced the pathway-specific calorimetric yields *β* and *δ*, which relate the outcome of metabolic activity in the form of either biomass or maximal growth rate to the reaction enthalpies *H*_+_ and *H*_−_, respectively. Due to the lack of metabolic adaptability of JCVI-Syn3B, we assumed that the molar reaction enthalpies of catabolic and anabolic reactions were not affected by the lipidome. Our data confirm this assumption, because the biomass-related heat release stayed in proportion to the independently determined cell volume, i.e., the biomass yield *β* did not depend on the lipidome (Fig. 3A). In contrast, the maximal rate of biomass production (*m·r*_*0*_) by JCVI-Syn3B cells, which we determined on a relative scale, could only be reproduced with a lipidome-dependent yield *δ* for the cell division rate (Fig. 4C). The efficiency of cell division decreased with biomass. The inverse (1/*δ-*) scales with the heat that is released to support a cell division rate of 1 h^−1^ and is thus a measure of the entropic cost *σ* of cell division (Table S1). The negative correlation between biomass and the entropic cost of maintaining high division rates indicates that both processes compete for the same energy source, which we believe is the shared cellular ATP/GTP pool. Each ATP molecule diverted from anabolism to the cell division machinery reduces biomass and increases the energy flux in the cell division pathway roughly proportionally. This relation is consistent with the almost constant *ΔH*_+_*/Δ* ratio, implying that the underlying biochemical reaction networks remained mostly unaltered while their relative energy consumption varied in the diets. As a consequence, *H*_+_ and *H*_–_ trace out an almost straight line in the “fitness landscape” in Fig. 4(B). An analogous competition between biomass production and glycolysis-generated ATP was previously observed in anaerobically fermenting yeast.(26) In this case the diversion of one ATP out of the four molecules produced during glycolysis reduced the biomass by 25%. In JCVI-Syn3B cells, the same competition for a shared glycolysis-generated ATP pool prevails, but our data indicate that it is the increased entropic cost for futile ATP hydrolysis during non-optimal lipidome-divisome interactions which creates a sink for ATP against which biomass formation has to compete. Thus, we identify the increased entropic cost *σ* of cell division as the major determinant of lipidome-dependent increase of metabolic heat flow.

What mechanisms could underlie the lipidome-dependent variation in energy dissipation during cell division in JCVI-Syn3B cells? Lipids influence membrane function through both bulk biophysical properties and chemically specific interactions with membrane-associated proteins. Protein-lipid interactions can involve hydrophobic contacts within transmembrane domains, as well as electrostatic and hydrogen-bonding interactions with peripheral membrane proteins. Changes in bulk membrane properties could, in principle, alter cellular energy efficiency: increased membrane permeability necessitates greater energy expenditure to maintain essential electrochemical gradients, while increased membrane rigidity could raise the energetic cost of membrane deformation during cell division. However, based on measurements of lipid order performed in parallel experiments(18), we can rule out membrane permeability as primary drivers of the observed metabolic inefficiency. Lipid order typically correlates with membrane permeability and bending rigidity — higher order corresponds to lower permeability and increased stiffness(27). The 2FA-PC diet had the lowest lipid order (highest permeability) but showed only marginal “extra” metabolic heat flow and displayed the maximal division rate while still achieving 80% of maximal biomass formation. These findings strongly suggest that variations in bulk membrane properties alone do not explain the lipidome-dependent differences in heat dissipation, thus implicating non-optimal lipid-protein interactions as the underlying cause.

We had initially hypothesized that the lipidome may primarily affect the efficiency of membrane-bound transporters for glucose and other nutrients through altered lipid-protein interactions, thereby reducing biomass accumulation. However, this was not supported by our data since we did not observe any correlation of lipidome-induced changes in nutrient flux per cell surface with heat dissipation or division rate (Fig. 2). Instead, variation of the energy efficiency of cell division was implicated by the relationship between growth rate, cell size and biomass production. The divisome relies on interactions of the lipid bilayer with integral membrane proteins and membrane-associated proteins required for force generation during cell division. Our data show that the entropic cost of cell division was high for plasma membranes that contained large fractions of negatively curved and/or negatively charged lipids (Fig. 4(C)) indicating that intrinsic membrane curvature and lipid net charge are key factors for divisome energetics. During cell division, CL may be favorably attracted by plasma membrane invaginations via curvature-induced lipid sorting.(28) The transition through differently shaped membrane structures may be one reason why lipid diversity, rather than a single “optimal” lipid class correlates with large CGP values. Similarly, the assembly of the proteins of the divisome and the attachment to the polymerizing FtsZ monomers in the cytosol strongly depends on physical lipid properties as shown for the headgroup charge-dependent rearrangement of ZipA-tethered FtsZ filaments on supported bilayers.(29) A plasma membrane of diverse lipid composition can provide the required charge and curvature contrast to allow localization of membrane anchored force-generating structures. Without such contrast, the attachment site may diffuse at the plasma membrane surface which would impair stable directionality during daughter cell formation. This appears physically reasonable, because the extracellular entropy increase by enhanced metabolic heat is the energetic equivalent of the maximal structure-generating potential (intracellular entropy decrease) which can be used to repeatedly rebuild force-generating structures with impaired membrane anchorage. We propose that inefficient FtsZ-membrane interactions generate a drain of ATP away from biomass synthesis to an impaired divisome machinery. This explanation finds an interesting parallel in minimal cell evolution. It has been shown that FtsZ evolution is directly reflected in the variation of cell sizes.(30) While in our study a “stable FtsZ” may tune cell size in response to lipidome variation, an “evolutionarily dynamic FtsZ” population does the same in response to FtsZ mutation. Both assays seem to address the same target, i.e., the efficiency of FtsZ-membrane interactions, from the side of either the lipidome or the protein.

We have shown that the complex interplay between metabolism, growth and cell division is well reproduced by a flux-model in which the lipidome affects only two components: the activation barriers of conformational transitions required for divisome function and the energy efficiency of the divisome. Figure 5C shows three representative situations which result from the model. A minimal lipidome impairs membrane anchorage and force generation of the divisome. Additional GTP cycles for reassembly of the membrane-bound components make up for the impairment at the expense of cell biomass but with little effect on intrinsic divisome kinetics (2FAs diet). A diverse and fluid lipidome supports optimal function of the divisome at high rate and energy efficiency leading to relatively high cell mass at high growth rate (4FAs_PC_SM). Finally, a plasma membrane of diverse composition with respect to charge and intrinsic lipid curvature supports an energy-efficient cell division but prolongs the doubling time by increased energy barriers, thereby allowing more biomass production between cell divisions (2FAs_SM diet).

## Conculsions

Key to the analysis of metabolic heat flow from JCVI-Syn3B cells was the initially heuristic use of the Monod equation to reach at an estimate of the biomass-related reaction enthalpy in relation to a reference culture. We think that the simplicity of the metabolism of JCVI-Syn3B cells was crucial for further developing the heuristics into a physically consistent simulation of the relations between metabolism, growth and cell division based on their calorimetric signatures. This emphasizes the enormous value of genomically streamlined cellular models for the analysis of fundamental physical properties of metabolism. In the context of previous enthalpy-based analyses of IMC data, the current work has helped developing the public software “metabolator” (metabolator.hzdr.de) for the evaluation of microbial heat flow curves.

## Materials and Methods

### Bacterial growth medium

The JCVI-Syn3B bacterial strain was received from J. Craig Venter Institute (JCVI) (La Jolla, California, USA)Chemicals and cultivated in SP4 growth medium, supplemented with CMRL. The original SP4 recipe was first developed by Tully et.al(31) and later supplemented with additional nutrients to support viability of the minimal cell.(16) The SP4 composition is listed in Table 1.

**Table 1.** SP4 growth medium composition (per 1L) for JCVI-Syn3B cultures.

| SP4 growth medium, pH 7.0 |  |  |
| --- | --- | --- |
| Component | Amount | Company/Details |
| Difco PPLO Broth | 3.5 g | Difco/Per 1L:<br>Beef heart infusion from 50 g – 6 g<br>Peptone – 10 g<br>NaCl – 5 g |
| Tryptone (peptone from casein) | 10 g | Sigma Aldrich |
| Peptone (peptone from gelatin) | 5.3 g | Sigma Aldrich |
| 20% yeastolate | 10 ml | Gibco/Bacto TC yeastolate |
| 15% yeast extract | 35 ml | Carl Roth |
| 7.5% NaHCO<sub>3</sub> | 14.6 ml | Honeywell |
| 20% glucose | 25 ml | Carl Roth |
| 400000 U/ml penicillin G, Na-salt | 2.5 ml | Carl Roth/Cellpure > 1550 U/mg, Penicillin G Na-salt |
| 25 mg/ml L-glutamine | 5 ml | Carl Roth/> 99% purity, Cellpure |
| CMRL | 3.92 g | HIMEDIA, CMRL 1066 Medium, w/o L-Glutamine, phenol red and sodium bicarbonate |
| Sodium bicarbonate | 0.88 g | Honeywell |
| Add H<sub>2</sub>O to the final volume of 950 ml |  |  |

### Monitoring metabolic activity by isothermal microcalorimetry (IMC)

JCVI-Syn3B cells were adapted to grow with each lipid diet over three passages (extended to five passages for 2FAs and 4FAs) prior the experiment. Cells were passaged daily to exclude ageing effects on growth. Batch bacterial cultures were setup in 100 ml glass flasks. Cells were harvested at mid-exponential phase and diluted in their native growth medium (SP4 + cyclodextrin-lipid complexes) to the starting OD 600 nm = 0.005. 2 ml were transferred to autoclaved calorimetric 4 ml glass ampoules (Waters GmbH, Eschborn, Germany), sealed manually with disposable tin lids. IMC was performed with a Thermal Activity Monitor, TAMIII (Waters GmbH, Eschborn, Germany) equipped with 12 single microcalorimeter channels which recorded the metabolic heat flow produced by the bacterial cultures continuously at 37°C for up to 100 h. Before measurement, the samples were held in the TAM III in a waiting position for 15 minutes before complete insertion in the microcalorimeter followed by additional 45 minutes of equilibration. Thermal changes in each ampoule were recorded as a continuous electronic signal (in Watts), which is proportional to the heat production rate.

Time-dependent heat flow curves were analysed as described earlier.(20) The data were first expressed as a function of the released heat (enthalpy plot) and then fitted with a hyperbolic dependence on nutrient concentration for the metabolic activity per cell. The mathematical approach is equivalent to applying the Monod equation over the largest possible data range in which the standard deviation of the fit from the raw data stayed below one percent of the heat flow maximum. By this criterion, the raw data covered 75% to 85 % of the predicted total heat release. The procedure determines three parameters to best reproduce the enthalpy plot of the raw data: *r*_*0*_: maximal division rate per cell; *H*_*0*_: extrapolated released heat of a culture grown to full consumption of accessible nutrient; *H*_*f*_: the heat released from a culture containing a nutrient concentration that supports initial growth at half-maximal rate up to full consumption of all accessible nutrient (see SI for more details of heat flow analyses).

### Cell size and cell number determinations

#### a)Dynamic light scattering

JCVI-Syn3B cells were grown to mid-exponential stage at 37°C with all lipid diets in biological triplicates in batch cultures. 7 ml of cell culture was harvested and centrifuged to remove medium (9000g, 2 min). Cells were resuspended in 1 ml of preheated MWB (200 mM NaCl, 25 mM HEPES, 1% glucose, pH 7.0) and the absorbance of the cell suspension determined at OD600 nm using DeNovix DS-11 FX+ spectrophotometer. Cells were diluted to OD 0.1 in MWB and incubated at 37°c for 5 min, 500 rpm shaking. The suspension was transferred to a 1 ml disposable cuvette (DTS0012) and its precise concentration (OD 600) was again measured. The average cell diameter of cells in MWB was determined by Malvern Zetasizer, with the following settings:

|  |  |
| --- | --- |
| Material | Liposomes |
| RI (material) | 1.45 |
| Absorption | 0.001 |
| Dispersant | Water |
| Temperature | 37.0°C |
| Viscosity | 0.6864 cP |
| RI (dispersant) | 1.33 |

The instrument was preheated to 37°C and the sample cuvette equilibrated for 10 min. 3 consecutive measurements were performed on the same cuvette, yielding total 9 measurements from 3 biological replicates. The count rate for JCVI-Syn3B samples was obtained simultaneously with cell size measurements. The built-in calculator tool in Malvern Zetasizer software allows for the total particle number in the sample estimation, based on the derived count rate (kilo-particles per second) and particle radius (nm). Resulting total number of particles/cuvette was normalized for sample OD 600 to obtain particle/ODU count.

#### b)Spot assays

SP4+CMRL+FBS agar plates were prepared as follows: 1.5% agar was mixed in with Basal SP4 and autoclaved; the mixture was let to cool down to 60°C and kept in 55°C water bath afterwards. In the meantime, the rest of SP4 components were mixed together, filter sterilized and heated to 37°C. The two media parts were quickly mixed together with a Pasteur pipette to yield a homogenous solution with no air bubbles. 25 mL agar-SP4 mixture were added to petri dishes and let solidify under the safety cabinet. The plates were subsequently exposed to UV light for 30 min. Prepared plates were stored at 4°C. JCVI-Syn3B cells were grown in batch cultures in triplicates at 37°C to mid-exponential and their density was determined using OD600 absorbance. Cell suspension was diluted in native SP4 + cyclodextrin + lipids to final OD 600 = 0.1. The precise cell absorbance of diluted samples was measured once again. Cell suspensions were serially diluted in their native growth medium in 96-well plates, with up to 10^−10^ dilution. 5 µL of diluted samples were seeded on preheated agar plates, using multichannel pipettes. From each biological replicate, 2 identical rows were seeded per plate, yielding total 6 spots for each dilution. Plates were let under the safety hood to dry, then sealed and kept at 37 °C in the incubator and images taken after 7-9 days.

CFUs (colony-forming-units) were manually counted for all spots, displaying the individual colonies. For very diluted spots, the colonies were counted manually, for intermediate dilutions that showed individual yet too numerous colonies, the Colony Counter plugin for Image J was used to determine the number of individual colonies. Finally, colony counts were renormalized back to the original cell sample absorbance to obtain CFU/ODU. The final value is calculated as the average CFU count per spot per plate.

## References

1. M. Woubshete, S. Cioccolo, B. Byrne, Advances in Membrane Mimetic Systems for Manipulation and Analysis of Membrane Proteins: Detergents, Polymers, Lipids and Scaffolds. Chempluschem 89 (2024).

2. I. G. Denisov, S. G. Sligar, Nanodiscs for the study of membrane proteins. Curr Opin Struc Biol 87 (2024).

3. H. Ayub et al., GPCRs in the round: SMA-like copolymers and SMALPs as a platform for investigating GPCRs. Arch Biochem Biophys 754 (2024).

4. J. Sarkis, V. Vié, Biomimetic Models to Investigate Membrane Biophysics Affecting Lipid-Protein Interaction. Front Bioeng Biotech 8 (2020).

5. E. Fischermeier et al., Dipolar Relaxation Dynamics at the Active Site of an ATPase Regulated by Membrane Lateral Pressure. Angewandte Chemie-International Edition 56, 1269–1272 (2017).

6. M. Doktorova et al., Cell membranes sustain phospholipid imbalance via cholesterol asymmetry. Cell 188, 2586–2602 e2524 (2025).

7. J. R. Winnikoff et al., Homeocurvature adaptation of phospholipids to pressure in deep-sea invertebrates. Science 384, 1482–1488 (2024).

8. D. H. Johnson, O. H. Kou, N. Bouzos, W. F. Zeno, Protein-membrane interactions: sensing and generating curvature. Trends Biochem Sci 49, 401–416 (2024).

9. S. Arumugam, P. Bassereau, Membrane nanodomains: contribution of curvature and interaction with proteins and cytoskeleton. Essays Biochem 57, 109–119 (2015).

10. C. Klose et al., Flexibility of a eukaryotic lipidome--insights from yeast lipidomics. PLoS One 7, e35063 (2012).

11. G. Chwastek et al., Principles of Membrane Adaptation Revealed through Environmentally Induced Bacterial Lipidome Remodeling. Cell Rep 32, 108165 (2020).

12. R. M. Wagner, L. Kricks, D. Lopez, Functional Membrane Microdomains Organize Signaling Networks in Bacteria. J Membrane Biol 250, 367–378 (2017).

13. A. Musatov, E. Sedlák, Role of cardiolipin in stability of integral membrane proteins. Biochimie 142, 102–111 (2017).

14. I. Justice, P. Kiesel, N. Safronova, A. von Appen, J. P. Saenz, A tuneable minimal cell membrane reveals that two lipid species suffice for life. Nat Commun 15, 9679 (2024).

15. C. A. Hutchison et al., Design and synthesis of a minimal bacterial genome. Science 351 (2016).

16. D. G. Gibson et al., Creation of a Bacterial Cell Controlled by a Chemically Synthesized Genome. Science 329, 52–56 (2010).

17. N. Safronova, L. Junghans, J. P. Saenz, Temperature change elicits lipidome adaptation in the simple organisms Mycoplasma mycoides and JCVI-syn3B. Cell Rep 43, 114435 (2024).

18. N. Safronova, L. Junghans, J. Oertel, K. Fahmy, J. P. Saenz, Chemically defined lipid diets reveal the versatility of lipidome remodeling in genomically minimal cells. bioRxiv 10.1101/2024.10.04.616688, 2024.2010.2004.616688 (2024).

19. J. Oertel, S. Sachs, K. Flemming, M. H. Obeid, K. Fahmy, Distinct Effects of Chemical Toxicity and Radioactivity on Metabolic Heat of Cultured Cells Revealed by “Isotope-Editing”. Microorganisms 11 (2023).

20. K. Fahmy, Simple Growth-Metabolism Relations Are Revealed by Conserved Patterns of Heat Flow from Cultured Microorganisms. Microorganisms 10 (2022).

21. Y. Wang, H. L. Zhu, J. G. Feng, P. Neuzil, Recent advances of microcalorimetry for studying cellular metabolic heat. Trac-Trend Anal Chem 143 (2021).

22. D. Fessas, A. Schiraldi, Isothermal calorimetry and microbial growth: beyond modeling. J Therm Anal Calorim 130, 567–572 (2017).

23. M. R. Tan et al., Detection of microorganisms in different growth states based on microcalorimetry. J Therm Anal Calorim 109, 1069–1075 (2012).

24. T. Maskow et al., What heat is telling us about microbial conversions in nature and technology: from chip-to megacalorimetry. Microb Biotechnol 3, 269–284 (2010).

25. Z. R. Thornburg et al., Fundamental behaviors emerge from simulations of a living minimal cell. Cell 185, 345–360 (2022).

26. A. C. A. van Aalst et al., Pathway engineering strategies for improved product yield in yeast-based industrial ethanol production. Syn Syst Biotechno 7, 554–566 (2022).

27. J. Steinkühler, E. Sezgin, I. Urbančič, C. Eggeling, R. Dimova, Mechanical properties of plasma membrane vesicles correlate with lipid order, viscosity and cell density. Communications Biology 2, 337 (2019).

28. E. Beltran-Heredia et al., Membrane curvature induces cardiolipin sorting. Commun Biol 2, 225 (2019).

29. P. Mateos-Gil et al., FtsZ polymers bound to lipid bilayers through ZipA form dynamic two dimensional networks. Biochim Biophys Acta 1818, 806–813 (2012).

30. R. Z. Moger-Reischer et al., Evolution of a minimal cell. Nature 620, 122–127 (2023).

31. J. G. Tully, D. L. Rose, R. F. Whitcomb, R. P. Wenzel, Enhanced Isolation of Mycoplasma-Pneumoniae from Throat Washings with a Newly Modified Culture-Medium. J Infect Dis 139, 478–482 (1979).

