## Supplementary Information for "Metabolic heat flow from the minimal cell JCVI-Syn3B reveals the lipidome-dependence of growth and metabolism"

### 1. Fundamental relations derived from the extended calorimetric Monod equation

#### 1.1 Shared and non-shared origins of metabolic heat flow from Mycoplasma cultures in different diets

The analysis of lipid-specific metabolic heat flow contributions is based on the comparison between the heat flow curve from a "sample" (*s*) diet, which releases the total heat $H_{0s}$, with that of a "reference" (r) diet. As reference, the 2FA_SM diet was chosen in which total heat dissipation was minimal, indicative of the most energy-efficient metabolism. All other heat flow curves were decomposed into a reference-like growth coupled to a released heat $H_{gs} = \alpha \cdot H_{0s}$ , representing the enthalpy contribution from the shared biomass-generating biochemical reactions. The additionally released "extra" metabolic heat $H_e = (1-\alpha) \cdot H_{0s}$ was then assigned to a lipididome-specific metabolic heat flow in the "sample diet".

This formal distribution of heat flow splits the "extended calorimetric Monod equation" (ECME) – the left side of Eq S1 – into two terms that are required to express the measured metabolic heat flow $P(H)_s$ as a function of cumulated released heat $H$. The first term uses the reference parameter $H_{fr}$ to model the nutrient dependence of normal growth-related metabolism. The second term complements the equation such that the sum of both terms is in accordance with the experimentally derived parameter $H_{fs}$ obtained for the "sample diet".

$$P(H)_s = r_{0s} \cdot H \cdot \frac{H_{0s} - H}{H_{0s} - H + H_{fs}} = H \cdot \left[ r_{0r} \cdot \frac{\alpha(H_{0s}-H)}{\alpha(H_{0s}-H)+H_{fr}} + \Delta r_0 \cdot \frac{[1-\alpha]\cdot(H_{0s}-H)}{[1-\alpha]\cdot(H_{0s}-H)+\Delta H_f} \right] \qquad \text{Eq S1}$$

with $\Delta H_f = H_{fs} - H_{fr},\ \Delta r_0 = r_{0s} - r_{0r},\ \alpha = H_{fr}/H_{fs}.$

Eq S1 shows that the decomposition of the measured heat flow curve into two metabolic pathway-specific hyperbolic nutrient dependencies is achieved by defining the fraction $\alpha$ as the ratio of the $H_f$ of the Monod constants of growth in the two diets, see 1.2). Averaged values of the fit parameters were obtained from three independent fits of triplicate heat flow curves shown in Fig. S1, the raw data were not averaged. This heuristic dissection of metabolic activity into shared and non-shared biochemistry predicts a growth-related heat release of $H_g$ which provides a relative measure of the biomass produced in relation to the reference culture. In combination with the relative cell volumes $V_c$ obtained from DLS (scaled to the reference cell volume obtained for the 2FAs_SM diet), also the relative cell numbers can be extrapolated to complete nutrient consumption as : $N_{tot}=H_g/V_c$. With this additional information, also the relative yield of cell formation $Y_{NH} = N_{tot}/H_0$ in the various diets is obtained. This allows the determination of $p_0$, the relative maximal thermal power per cell, from the relation $r_0=p_0\cdot Y_{NH}$. Finally, these numbers can be used to reconstruct the measured heat flow curves in the time domain according to Eq S1:

$$P(t) = p_0 \cdot [N_0 + Y_{NH} \cdot H(t)] \cdot \frac{H_0 - H(t)}{H_0 - H(t) + H_f} \qquad \text{Eq S2}$$

with $N_0$ the inoculate size. Figures S2 A and B show that the yields $Y_{NH}$ correctly predict the time at which peak metabolic activity was reached as exemplified for two diets in which JCVI-syn3B cells grew faster or slower than the reference culture.

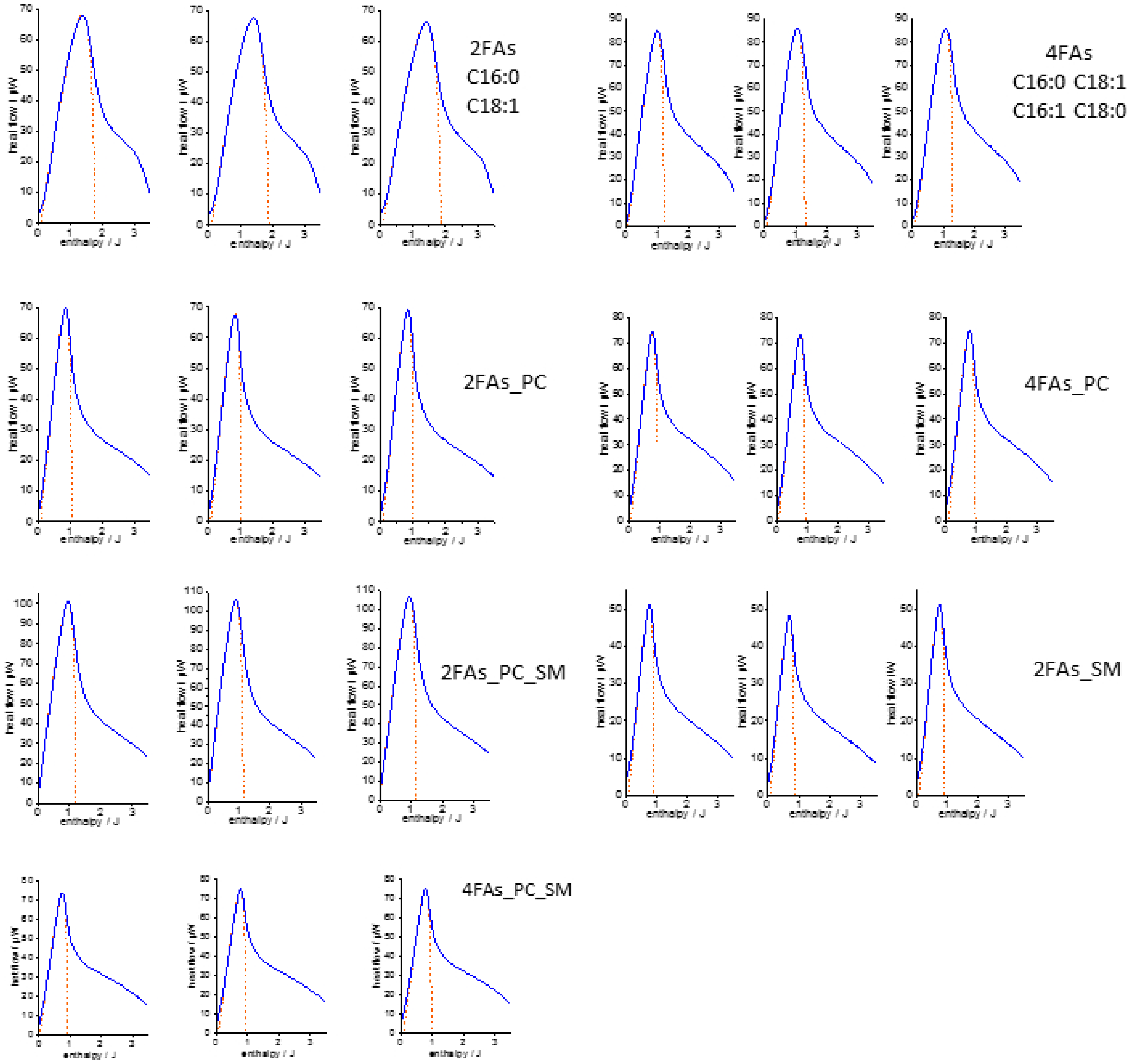

**Figure S1: Raw data of heat flow curves recorded in triplicates for growth of JVIsyn3B in the indicated diets.** All experiments were carried out with a TAMIII instrument (TA-waters, Eschborn) at 37 °C in 4 mL ampoules filled with 2 mL of defined growth medium containing the indicated lipid diets (see main text for details).

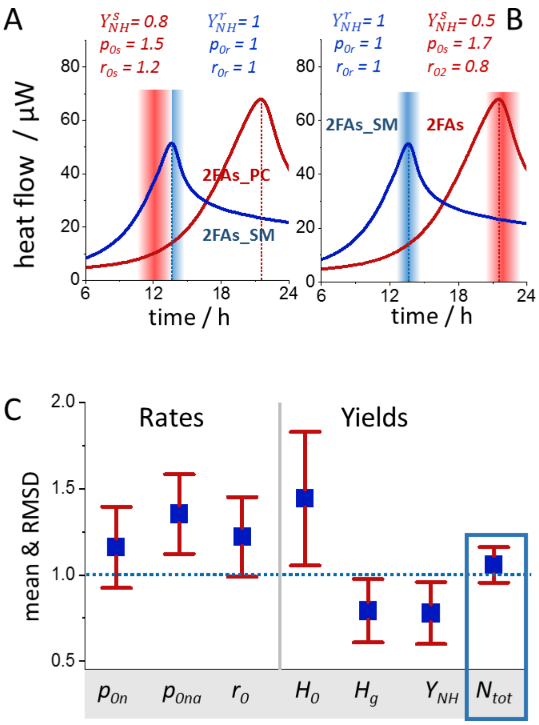

**Figure S2: Consistency of enthalpy-based derivations of rates and yields with the time-dependent heat flow curves.** The cell number in the inoculate of the reference culture was 1.4x10$^9$ (SD = 0.3x10$^9$) as determined by a CFU Spot assay from 12 measurements. This allowed determining $Y_{NH}^{r}$ from the exponential growth phase of the reference culture grown in the 2FAs_SM diet. Therefore, the maximal heat flow per cell $p_0$ could be obtained from the calorimetrically determined rates $r_0$ and the time-dependent heat flow curves reconstructed on an absolute time scale according to Eq S2 (From the enthalpy plots only a relative time scale is obtained with an initially undetermined integration constant which is chosen to match maximal heat flow peaks in the fitted and measured data). A) comparison of the predicted peak times for growth in the reference culture (blue) and the 2FAs_PC diet (red). The colored bars show the predicted peak occurrence with a jitter of about 2.5 h, due to handling and equilibration time before heat flow recordings. B) The same comparison for the 2FAs diet. C) Statistics of all determined yields and rates (including the reference diet). Dotted line corresponds to the parameters of the reference culture set to unity.

### 1.2 Relation between $H_f$ and the classical Monod constant

The Monod term in calorimetric representation can be formally transformed into the classical concentration-based Monod constant according to:

$$\frac{H_0 - H(t)}{H_0 - H(t) + H_f} = \frac{Y_{SH}^{v}(H_0 - H(t))}{Y_{SH}^{v}(H_0 - H(t) + H_f)} \qquad \text{Eq S3}$$

with $Y_{SH}^{v}$ the yield coefficient which gives the amount of metabolized substrate in proportion to released metabolic heat. The superscript $v$ indicates further division by the sample volume, such that the product of $Y_{SH}^{v}$ with heat is a substrate concentration. All cultures were grown in the same nutrient concentration and volumes such that their $H_0$ values correspond to the same amount of consumed substrates. Therefore, the yield coefficient is easily determined as $Y_{SH}^{v} = [S_0]/H_0$ and the Monod constant is

$$C_M = [S_0] \cdot \frac{H_f}{H_0} = Y_{SH}^{v} \cdot H_f \qquad \text{Eq S4}$$

A single rate-limiting nutrient component is not known and the $C_M$ values in Table I are given formally for glucose which represents the main energy and carbon source with $[S_0]$= 50 g L$^{-1}$.

With the metabolic dissection presented in 1.1 another fundamental property follows for the interpretation of the $H_f$ and $C_M$ values. Taking the reference culture as the standard of biomass production with a yield $Y_{BH}^{r}$ (in g per Joule) the biomass formation $B_{ms}$ of a sample diet is related to the biomass yield of the reference by:

$B_{ms} = Y_{BH}^{r} \cdot \alpha \cdot H_{0s}$. (Since the biomass was too small to be determined, the absolute value of $Y_{BH}^{r}$ is not known and the heat equivalent $H_g = \frac{H_{fr}}{H_{fs}} \cdot H_{0s}$ is used as a proxy of biomass in the main text).

The relative biomass is: $\frac{B_{ms}}{B_{mr}} = \frac{H_{fr}}{H_{fs}}\frac{H_{0s}}{H_{0r}}$, which by definition equals $\frac{Y_{BH}^{s} \cdot H_{0s}}{Y_{BH}^{r} \cdot H_{0r}}$.

Thus: $$H_{fr} \cdot Y_{BH}^{r} = H_{fs} \cdot Y_{BH}^{s} = b_m \qquad \text{Eq S5}$$

This shows that in all diets, the same amount of biomass $b_m$ is formed when the cultures are grown with an initial amount of nutrient corresponding to their individual metabolic heat equivalents $H_f$. With Eq S4 follows for the Monod constants:

$$C_M = \frac{b_m \cdot [S_0]}{Y_{BH} \cdot H_0} = \frac{b_m \cdot [S_0]}{B_m} \qquad \text{Eq S6}$$

Since $b_m$ and $[S_0]$ are constants for all diets, the Monod constants are inversely related to the total biomass $B_m$ formed in each culture. The linear relation between $H_g$ and cell volume, derived from experiment, proves that the calorimetrically determined $C_M$ values are in fact consistent with expressing biomass in proportion to $H_g$. This further reveals that the total lipidome-dependent biomass formation in the diets is predominantly attributable to cell volume, rather than to effects on cell numbers (Fig. S1).

#### 1.3 Correlations between rates and yields

##### 1.3.1 Biomass yield expressions

We express the reference-based biomass yield $Y_{BH}^{r}$ (biomass formed per heat release in relation to the reference culture) by

$$Y_{BH}^{r} = \frac{B}{H_0} = w \cdot \frac{H_g}{H_0} \qquad \text{Eq S7}$$

*B* is the total biomass of a culture and $\omega$ is the expressing the biomass produced per heat release by the reference diet. The underlying biochemistry is shared among the cells in all diets, rendering $\omega$ independent of the lipidome. Thus, $\frac{H_g}{H_0}$ is a relative measure of the "true" diet-specific biomass yield $Y_{BH}$. According to the definition, $H_e$ can be used to express, $\frac{H_g}{H_0}$ such that:

$$Y_{BH}^{r} = w \cdot \frac{\frac{H_g}{H_e}}{1+\frac{H_g}{H_e}} = w \cdot \frac{\frac{s_g}{s_e}}{v_r + \frac{s_g}{s_e}} \qquad \text{Eq S8}$$

Here, $s_g$ and $s_e$ designate the amount of substrate utilization in reactions that are shared or not shared with the reference culture, respectively. When both types of substrate utilizations adopt the ratio $v_r$, both reaction routes contribute equally to metabolic heat release ($H_g$=$H_e$). Eq S8 shows that the biomass yield $Y_{BH}^{r}$ follows a hyperbolic saturation behavior with respect to relative substrate utilization with $\omega$ the maximal biomass yield per heat release defined by the 2FAs_SM reference culture.

In order to express these relations independently of a reference culture, we use the terms $H_+$ and $H_-$ (replacing $H_g$ and $H_e$) to define the amounts of heat formed by purely biomass-generating biochemistry and non-biomass-related biochemistry, respectively. This distinction is necessary because the heat $H_g$, although a reliable relative measure of biomass formation, it contains both types of enthalpic contributions and is thus larger than the purely biomass-related heat production. Consequently, the reference culture always achieves the maximal yield $Y_{BH}^r = w$, which is an unjustified physiological assumption. The true biomass yield is given without restriction of generality as:

$$Y_{BH} = \beta \cdot \frac{\frac{H_+}{H_-}}{1+\frac{H_+}{H_-}} = \beta \cdot \frac{\frac{s_+}{s_-}}{v+\frac{s_+}{s_-}} = \beta \cdot \frac{e^{ln\frac{s_+}{s_-}-\ln v}}{1+e^{ln\frac{s_+}{s_-}-\ln v}} \quad \text{Eq S8a}$$

with $H_+ < H_g$ and $H_- > H_e$. In contrast to the overall yield $Y_{BH}$, the pathway-specific yield parameter $\beta$ expresses biomass as $m = \beta \cdot H_+$. Eq S8a shows that $Y_{BH}$ follows a typical saturation curve with respect to the substrate utilization ratio $\frac{s_+}{s_-}$, i.e, a logistic curve with respect to $ln\frac{s_+}{s_-}$. For $\frac{s_+}{s_-} = v$, the half-maximal biomass yield of the metabolic network is reached. The reference-based and reference-independent heats are related through: $H_+ = H_g - H_g \cdot f_-^r$ and $H_- = H_e + H_g \cdot f_-^r$ , with $f_-^r$ the fraction of non-biomass-related heat production in the reference culture. A factor $f_-^r = 0.17$ was required to correct the reference-based yields such that they followed the normalized logistic function. Plotted *vs* the same variable $ln\frac{H_+}{H_-}$, the calorimetrically determined maximal division rates adopt a logistic distribution. The parameter $f_-^r$ rescales the reference-based biomass proxy $\frac{H_g}{H_0}$ by the factor $(1 - f_-^r)$ such that the reference culture exhibits a true biomass yield of $Y_{BH} = 0.83$ on the correctly normalized scale between zero and unity (Fig. 4(A) in the main text).

#### 1.3.2 Rate expressions

By definition, the rate of per cell biomass production is given by:

$$\frac{dB}{dt} = p_0 \cdot Y_{BH} = m_c \cdot r_0 \quad \text{Eq S9}$$

with $m_c$ the cellular biomass. With $p_{0m} = \frac{p_0}{m}$ , the specific heat flow (heat flow per cell mass), it follows:

$$r_0 = p_0^m \cdot Y_{BH} \quad \text{Eq S9a}$$

The bell-shaped dependence of $r_0$ on that the specific heat flow $p_{0m}$ is not constant, because the plot of the rates against $ln\frac{H_+}{s_-}$ did not follow the sigmoidal shape of the biomass yield $Y_{BH}$ (Fig. 4(A)). Instead, our data show that the rates are only correctly reproduced, when the specific heat flow is described by a negative linear dependence on $Y_{BH}$:

$$p_{0m} = P_0^m \cdot (1 - \frac{Y_{BH}}{\beta}) \quad \text{Eq S10}$$

where $P_0^m$ is a constant describing the limiting maximal specific heat flow, (theoretically achieved when $Y_{BH} = 0$, i.e., metabolic activity would exclusively be used for non-biomass processes). Thus, the specific thermal metabolic power $p_{0m}$ decreases with increasing biomass yield as seen already for the plot against $Y_g$ (Fig. 3(B)). The experimentally obtained values of maximal specific heat flow $\boldsymbol{p_{0m}}$ for the different diets shown in Fig. 3(D) was modeled according to Eq S10 with $P_0^m = 4.6$ and $\beta = 1.06$.

Inserting $p_{0m}$ in Eq S9a leads to the experimentally observed parabolic dependence of $r_0$ on $Y_{BH}$ :

$$r_0 = P_0^m \cdot (1 - \frac{Y_{BH}}{\beta}) \cdot Y_{BH} \qquad \text{Eq S10a}$$

The yield $Y_{BH}$ follows a logistic curve with respect to $ln\frac{s_+}{s_-}$ (Eq S8a and Fig. 4(B)). The number value of the latter expression is identical to the chemical potential difference between substrate components used for either biomass or non-biomass biochemistry. The term $(1 - \frac{Y_{BH}}{w}) \cdot Y_{BH}$ in Eq S10a equals the derivative of this logistic curve, i.e., it follows a logistic distribution. Accordingly, the most concise form of expressing the cell division rate of "minimal cells" as a function of the biomass yield is given by:

$$r_0 = P_0^m \cdot \frac{dY_{BH}}{d\Delta\mu} \qquad \text{Eq S11}$$

The division rates of minimal cells thus correlate with the sensitivity of the biomass yield to the change in the chemical potential difference $\Delta\mu$ of substrate used for biomass and non biomass-related metabolism.

Eq S10a is a special solution of a general growth relation. This becomes clearer when division rate $r_0$ is expressed in analogy to biomass (Eq S8a) by a pathway-specific yield parameter $\delta$ such that the division rate is $r_0 = \delta \cdot H_-$. Here, $\delta$ is the division rate per heat $H_-$ released from all non-biomass-producing metabolic processes that have occurred between two cell divisions. With this definition, the overall yield parameter (sum of all metabolic reaction enthalpies) of cell division rate can generally be expressed by:

$$Y_{DH} = \delta \cdot (1 - \frac{Y_{BH}}{\beta}) \qquad \text{Eq S12}$$

and

$$r_0 = Y_{DH} \cdot H_0 = \delta \cdot (1 - \frac{Y_{BH}}{\beta}) \cdot H_0 \qquad \text{Eq 12a}$$

Equating this with Eq 10a shows that for minimal cells:

$$\delta = P_0^m \cdot \frac{Y_{BH}}{H_0} \qquad \text{Eq S13}$$

and

$$\sigma = \delta^{-1} = \frac{H_0}{Y_{BH} \cdot P_0^m} \qquad \text{Eq S13a}$$

The inverse of $\delta$ measures the heat released per division rate, i.e., the entropic cost $\sigma$ of performing one cell division per hour (for the amount of biomass generated from the available nutrient).

Finally, the general calorimetric equation for the rate of biomass production adopts a symmetric form with the two introduced pathway-related total yields $Y_{BH}$ and $Y_{DH}$:

$$m \cdot r_0 = H_0 \cdot Y_{BH} \cdot H_0 \cdot Y_{DH} = \beta \cdot H_+ \cdot \delta \cdot H_- \qquad \text{Eq S14}$$

Here, $m$ is the extrapolated total biomass formed upon consumption of the initially available amount of nutrients. This equation allows comparing relations between biomass and cell division rate as a function of the pathway-specific yield parameters $\beta$ and $\delta$ as shown in Fig. 4(B). The "fitness landscape" in Fig. 4(B) in relative scale was produced by Eq S14 using $\beta = 1$, $\delta$ according to Eq S13 with $P_0^m = 1$.

**2. Lipids**

Cholesterol, palmitic acid (C16:0), oleic acid (C18:1), palmitoleic acid (C16:1) (Sigma), stearic acid (C18:0, Merck), 1-palmitoyl-2-oleoyl-glycero-3-phosphocholine (POPC) and egg sphingomyelin (SM) were purchased in pure form (Avanti) and stored at -20°C prior use. For working lipid stocks, POPC and SM were dissolved in pure EtOH to 100 mg/ml concentration. The precise mM concentration was further confirmed with phosphate assay. Cholesterol and fatty acids were dissolved in chloroform to yield concentrated stocks of the following concentrations:

| Lipid | Concentration, mM |
|---|---|
| cholesterol | 206.8 mM |
| C16:0 | 156 mM |
| C18:1 | 169.6 mM |
| C16:1 | 156 mM |
| C18:0 | 156 mM |

The chloroform stocks were kept in sealed glass vials at -20°C. Combined working EtOH stocks with cholesterol and fatty acids were prepared as follows:

| 2FAs stock | Concentration, mM | 4FAs stock | Concentration, mM |
|---|---|---|---|
| cholesterol | 51.7 mM | cholesterol | 51.7 mM |
| C16:0 | 78 mM | C16:0 | 39 mM |
| C18:1 | 81.4 mM | C18:1 | 42.4 mM |
| | | C16:1 | 39 mM |
| | | C18:0 | 39 mM |

The total lipid concentration was 211.1 mM. Lipids in chloroform were combined in 2 ml solvent-proof Eppendorf tubes and the solvent was evaporated using vacuum concentrator for 40 min. Next, 1 ml of pure EtOH (Uvasol) was added to the lipid film and the lipid stocks were incubated at 50°C for 20 min to ensure complete solubility of the lipid film. Additional lipid diets were generated from the above described 2FAs and 4FAs stocks designated 2FAs_PC and 4FAs_PC when supplemented with POPC; 2FAs_SM for supplementation with sphingomyelin; 2FAs_PC_SM and 4FAs_PC_SM when supplemented with both POPC and sphingomyelin.

**2.1 Cyclodextrin-lipid complexes and bacterial lipid feeding**

The cyclodextrins mβCD and mαCD were obtained from Sigma and CycloLab, respectively. 80 mM stocks of each cyclodextrin were dissolved in PBS (pH 7.0) and stored at 4°C. During SP4 growth medium preparation, the total $H_2O$ volume was reduced by 1/20 to leave space for cyclodextrin incorporation later: per 1 L total SP4, 950 ml stock was prepared.

2FAs and 4FAs stocks were complexed with mβCD, whereas mαCD was used for phospholipids. The different cyclodextrins were always incubated with their respective lipid types separately. For all cyclodextrin-lipid complexes, the lipid stock in EtOH was transferred to an Eppendorf tube and dried under vacuum. Cyclodextrin stocks were heated to 37°C and added to the dry lipid film. The samples were left overnight at 40°C with 1000 rpm shaking to ensure complete dissolution of dry lipid films.

Complexed stocks formed a homogenous, transparent solution and were stored at -20°C and used within one week. Stocks were heated to 37°C prior use and had the following concentration ratios in final SP4 for JCVI-Syn3B:

1.4 mM m$\beta$CD: 0.2 mM 2FAs/4FAs (for all diets)

2 mM m$\alpha$CD: 0.025 mM POPC/egg SM - for 2FAs-PC and 2FAs-SM diets

2 mM m$\alpha$CD: (0.0125 mM POPC + 0.0125 mM egg SM) - for (2FAs/4FAs-PC-SM) diets

Cyclodextrin-lipid complexes were prepared at 40x concentration to be diluted in SP4 medium. For final 10 mL of SP4 culture the recipe is as follows:

| | | | |
|---|---|---|---|
| For 2FAs/4FAs diets (no phospholipids) | | | |
| Final SP4 volume with lipids | Add SP4 stock | Add m$\beta$CD:FAs | Add PBS |
| 10 ml | 9.5 ml | 250 uL | 250 uL |
| For diet with FAs and phospholipids | | | |
| Final SP4 volume with lipids | Add SP4 stock | Add m$\beta$CD:FAs | Add m$\alpha$CD:PLs |
| 10 ml | 9.5 ml | 250 uL | 250 uL |

To prevent lipid precipitation in the growth medium, SP4 was warmed to 37°C prior to adding cyclodextrin-lipid stocks.

**3. Determination of membrane lipidome composition using shotgun lipidomics**

Bacterial samples were grown in batch cultures (in triplicates) at 37°C and harvested at early-to-mid exponential growth stage (Syn3B: OD 600 0.07-0.13) and washed twice in Mycoplasma wash buffer (200 mM NaCl, 25 mM HEPES, 1% glucose, pH 7.0) to remove traces of the growth medium at their growth temperature (21000g, 2 min centrifuging steps). A respective blank with SP4 + lipids was incubated together with bacterial cultures, washed and processed in the same way as the bacterial samples. Washed cell pellets and blank samples were resuspended in 150 uL PBS and flash-frozen in liquid nitrogen.

In brief, lipids were extracted using a two-step chloroform/methanol procedure.(1) Samples were spiked with internal lipid standard mixture containing: cardiolipin 14:0/14:0/14:0/14:0 (CL), ceramide 18:1;2/17:0 (Cer), diacylglycerol 17:0/17:0 (DAG), hexosylceramide 18:1;2/12:0 (HexCer), lyso-phosphatidate 17:0 (LPA), lyso-phosphatidylcholine 12:0 (LPC), lyso-phosphatidylethanolamine 17:1 (LPE), lyso-phosphatidylglycerol 17:1 (LPG), lyso-phosphatidylinositol 17:1 (LPI), lyso-phosphatidylserine 17:1 (LPS), phosphatidate 17:0/17:0 (PA), phosphatidylcholine 17:0/17:0 (PC), phosphatidylethanolamine 17:0/17:0 (PE), phosphatidylglycerol 17:0/17:0 (PG), phosphatidylinositol 16:0/16:0 (PI), phosphatidylserine 17:0/17:0 (PS), cholesterol ester 20:0 (CE), sphingomyelin 18:1;2/12:0;0 (SM), triacylglycerol 17:0/17:0/17:0 (TAG) and cholesterol D6 (Chol). After extraction, the organic phase was transferred to an infusion plate and dried in a speed vacuum concentrator. 1st step dry extract was re-suspended in 7.5 mM ammonium acetate in chloroform/methanol/propanol (1:2:4, V:V:V) and 2nd step dry extract in 33% ethanol solution of methylamine in chloroform/methanol (0.003:5:1; V:V:V). All liquid handling steps were performed using Hamilton Robotics STARlet robotic platform with the Anti Droplet Control feature for organic solvents pipetting.

Samples were analyzed by Lipotype GmbH (Dresden, Germany) using direct infusion on a QExactive mass spectrometer (Thermo Scientific) equipped with a TriVersa NanoMate ion source (Advion Biosciences). The data acquisition and quantitative evaluations have been described in detail.(2)

---

**Table S1:** Yield and rate parameters describing growth of JCVIsyn3B cells in various lipid diets. $p_0$: maximal heat flow per cell; $p_{0ga}$: maximal glucose uptake rate per µm$^2$ cell surface; $r_0$: maximal division rate per cell; $r_i$: initial growth rate; $H_0$: extrapolated released heat of a culture grown to full consumption of accessible nutrient; $H_f$: fitted of heat released from a culture growing initially at half-maximal rate up to full consumption of all accessible nutrient; $H_g$: growth related heat release upon full consumption of all accessible nutrient; $Y_{NH}$: number of cells formed per one pJ of metabolic heat release upon full consumption of all accessible nutrient; $C_M$: Monod constant; $D_c$: cell diameter; $N_{tot}$: extrapolated total number of cells formed upon consumption of all accessible nutrient (determined by CFU count for the 2FAs_SM diet, all other numbers expressed relative to the reference using the ratio $H_g/vol$). Relative and absolute values are given in red and blue print, respectively. All parameters are averages from three experiments.

*PG*: phosphoglycerol; PC: Phosphatidylcholine; SM: sphingomyelin; DAG: diacylglycerol; chol: cholesterol.

| Diets → parameters ↓ | 2FAs | 4FAs_PC | 4FAs | 2FAs_PC_SM | 2FAs_PC | 4FAs_PC_SM | 2FAs_SM |
|---|---|---|---|---|---|---|---|
| $p_0$ / fW | 1.67 | 2.29 | 1.42 | 2.16 | 1.48 | 1.43 | 1 |
| | *1.25* | *1.72* | *1.07* | *1.62* | *1.11* | *1.07* | *0.75* |
| $P_{0na}$ | 1.46 | 1.64 | 1.37 | 1.75 | 1.27 | 1.48 | 1 |
| $r_0$ / $h^{-1}$ | 0.80 | 1.37 | 1.24 | 1.52 | 1.19 | 1.41 | 1 |
| | *0.28* | *0.48* | *0.43* | *0.53* | *0.41* | *0.49* | *0.35* |
| $r_i$ / $h^{-1}$ | 0.76 | 1.33 | 1.21 | 1.52 | 1.21 | 1.39 | 1 |
| | *0.25* | *0.44* | *0.40* | *0.50* | *0.40* | *0.46* | *0.33* |
| $\delta$ / $h^{-1}$ $J^{-1}$ | 0.09 | 0.22 | 0.35 | 0.40 | 0.70 | 0.80 | 1 |
| $\sigma$ /h pJ | 11.47 | 4.56 | 2.85 | 2.52 | 1.43 | 1.25 | 1 |
| | *42.90* | *17.05* | *10.66* | *9.42* | *5.35* | *4.68* | *3.74* |
| $H_0$ / J | 2.14 | 1.83 | 1.47 | 1.43 | 1.16 | 1.06 | 1 |
| | *1.77* | *1.52* | *1.22* | *1.18* | *0.96* | *0.88* | *0.83* |
| $H_f$ / J | 6.00 | 1.50 | 2.25 | 2.00 | 1.50 | 1.25 | 1 |
| | *0.24* | *0.11* | *0.09* | *0.08* | *0.06* | *0.05* | *0.04* |
| $H_g$ / J | 0.40 | 0.73 | 0.75 | 0.81 | 0.94 | 0.91 | 1 |
| | *0.33* | *0.61* | *0.63* | *0.67* | *0.78* | *0.75* | *0.83* |
| $Y_{NH}$ /$pJ^{-1}$ | 0.48 | 0.60 | 0.87 | 0.71 | 0.81 | 1.00 | 1 |
| | *0.06* | *0.08* | *0.12* | *0.09* | *0.11* | *0.13* | *0.13* |
| $C_M$ / g $L^{-1}$ | 2.5 | 1.37 | 1.33 | 1.22 | 1.07 | 1.11 | 1 |
| | *0.67* | *0.37* | *0.36* | *0.33* | *0.29* | *0.30* | *0.27* |
| $D_c$ / nm | 0.73 | 0.85 | 0.84 | 0.93 | 1.00 | 0.95 | 1 |
| | *344* | *398* | *394* | *437* | *470* | *448* | *470* |
| $A_c$/$\mu m^2$ | 0.53 | 0.71 | 0.71 | 0.88 | 1.00 | 0.94 | 1 |
| | *0.09* | *0.12* | *0.12* | *0.15* | *0.17* | *0.16* | *0.17* |
| $V_c$ /$10^{-3}$ $\mu m^3$ | 0.39 | 0.66 | 0.59 | 0.80 | 1.00 | 0.87 | 1 |
| | *21.26* | *36.09* | *31.93* | *43.78* | *54.46* | *47.08* | *54.36* |
| $N_{tot}$ / $10^9$ | 1.03 | 1.10 | 1.28 | 1.01 | 0.94 | 1.05 | 1 |
| | *1.44* | *1.54* | *1.79* | *1.41* | *1.32* | *1.47* | *1.40* |
| CGP | 0.32 | 1.0 | 0.93 | 1.23 | 1.12 | 1.28 | 1 |
| % lipid classes | | | | | | | |
| PG | 27.6 | n.a. | 39.0 | 13.1 | 6.1 | 7.6 | 12.3 |
| PC | 0 | n.a. | 0 | 14.14 | 32.1 | 16.3 | 0 |
| CL | 16.9 | n.a. | 7.1 | 11.2 | 9.9 | 12.9 | 14.1 |
| DAG | 4.6 | n.a. | 0.6 | 5.9 | 0.4 | 6.7 | 1.6 |
| SM | 0 | n.a. | 0 | 8.1 | 0 | 10.8 | 18.5 |
| chol | 51.0 | n.a. | 53.0 | 47.6 | 51.5 | 45.7 | 53.4 |

**References**

1. C. S. Ejsing *et al.*, Global analysis of the yeast lipidome by quantitative shotgun mass spectrometry. *P Natl Acad Sci USA* **106**, 2136-2141 (2009).
2. N. Safronova, L. Junghans, J. Oertel, K. Fahmy, J. P. Saenz, Chemically defined lipid diets reveal the versatility of lipidome remodeling in genomically minimal cells. *bioRxiv* 10.1101/2024.10.04.616688, 2024.2010.2004.616688 (2024).
